# Toroidal displacement of *Klebsiella pneumoniae* by *Pseudomonas aeruginosa* is a unique mechanism to avoid competition for iron

**DOI:** 10.1101/2022.09.21.508880

**Authors:** Diana Pradhan, Ajay Tanwar, Srividhya Parthasarathy, Varsha Singh

## Abstract

Competition for resources is one of the major drivers for evolution and retention of new traits in microbial communities. Quorum-dependent traits of opportunistic human pathogen *Pseudomonas aeruginosa* allow it to survive and thrive in nature. Here, we report a unique surfactant-driven pushing mechanism that *P. aeruginosa* employs specifically against *Klebsiella pneumoniae*. The pushing is accomplished in a manner that is dependent on nutrient limitation and quorum sensing. We find that *P. aeruginosa* employs neither proteases nor toxic secondary metabolites against *K. pneumoniae*. Rhamnolipid biosurfactant appears to be the only factor required to displace Klebsiella effectively. Both rhamnolipid production and the pushing ability of *P. aeruginosa* are suppressed by iron supplementation. We show that both these bacteria produce several siderophores in minimal medium and rapidly deplete iron. Under these conditions, *P. aeruginosa* pushes Klebsiella away from the substratum using rhamnolipid, reducing the competition for iron. Our study describes a unique quorum and iron-responsive mechanism in *P. aeruginosa* to support its own growth during resource competition.

## INTRODUCTION

In most natural settings, microbes are generally known to occur as multispecies consortia including in pathophysiological conditions like cystic fibrosis, pneumonia, urinary tract infection, and diabetic foot ulcer. Microbes with diverse metabolic capacities can cooperate and coexist in a consortium. However, pathogens occupying a common niche might compete for macro or micronutrients resulting in altered virulence and persistence of one or more pathogens. These interactions are orchestrated inside a host or in the environment under resource-limiting conditions.

The behavior of a microbe can be profoundly altered by the presence of another microbe. Analyses of gene expression in specific pathogens show that expression pattern is altered in presence of another pathogen often leading to an increase in virulence of the former as reported for *Pseudomonas aeruginosa in vitro* (DeVault et al., 1990; Mashburn et al., 2005; Trejo-Hernández et al., 2014). *Staphylococcus aureus* induces the production of virulence factor, pyocyanin, in *P. aeruginosa* (Korgaonkar et al., 2013). Mono versus dual infection of the animal host also reflects an alteration in pathogenesis and disease progression (Hotterbeekx et al., 2017). Most human infections are polymicrobial (Brogden et al., 2005). Clinical outcomes for coinfections involving two or more pathogens are more severe than infection with a single pathogen (Marchaim et al., 2012; Okada et al., 2010). These include pneumonia (Cillóniz et al., 2011; Combes et al., 2002), wounds infection (Frank et al., 2009; Pastar et al., 2013), diabetic foot ulcer (Dowd, Wolcott, et al., 2008; Macdonald et al., 2021) and intestinal inflammation (Frisan, 2021). Therefore, there is a need to understand pathogenesis driven by inter-microbial interactions.

Bacterial pneumonia is often characterized by the presence of many bacteria. *P. aeruginosa, S. aureus, Klebsiella pneumoniae* and *Acinetobacter baumannii* are some of the commonly occurring antibiotic-resistant pathogens in hospital-acquired pneumonia (Jones, 2010). *P. aeruginosa* and *K. pneumonia* are known to share a common niche in cystic fibrosis lungs, in sepsis, and in chronic wound infections (Beaume et al., 2015; Dowd, Sun, et al., 2008; Rogers et al., 2003). Both species can coexist as a stable dual-species biofilm (Stewart et al., 1997). Under nutrient-replete conditions, LasB protease of *P. aeruginosa* is necessary for the dispersal of *in vitro* biofilm of *K. pneumoniae* (Childers et al., 2013). Intriguingly, *P. aeruginosa* growth is inhibited by *K. pneumoniae* in rich media whereas its growth is supported by *K. pneumoniae* under biotin limiting conditions via an unknown mechanism (Beaume et al., 2015). These reports show that the outcome of interactions between these two bacteria is highly contextual.

Pathogens with specific virulence factors have an advantage over neighboring bacteria. These factors include antibiotics and toxic molecules such as phenazines and hydrogen cyanide produced by *P. aeruginosa* (Borrero et al., 2014; Rijavec & Lapanje, 2016). Some bacteria also utilize specific surfactants to disperse from biofilm or use surfactant-aided motility such as swarming to claim resources in a given area (Caiazza et al., 2005; Davey et al., 2003). These behaviors are often orchestrated under nutrient-limiting conditions.

In this study, we examined environmental and molecular determinants of interactions between *P. aeruginosa* and *K. pneumoniae* under nutrient-limiting conditions. We established a coculture assay on agar surface and found that *P. aeruginosa* and *K. pneumoniae* coexist on complex, nutrient-rich media. On minimal media, *P. aeruginosa* rapidly pushes away *K. pneumoniae*. The displacement of *K. pneumoniae* by *P. aeruginosa* is independent of all its virulence factors except rhamnolipids. We find that the pushing behavior of *P. aeruginosa* can be suppressed by the supplementation of iron. On minimal media, *P. aeruginosa* and *K. pneumoniae* produce several siderophores each setting off competition for iron. This results in surfactant production in *P. aeruginosa* allowing it to push away non-motile *K. pneumoniae* from the surface. Our study has uncovered an additional function of the surfactant that bacteria *can* utilize to push competitors away.

## RESULTS

### Interaction of *P. aeruginosa* with *K. pneumoniae* is media dependent

To understand the rules of engagement between *P. aeruginosa* (Pa) and *K. pneumoniae* (Kp), we examined their interaction on solid media with varying nutritional content. We studied mono and coculture of Pa and Kp, 24 hours after plating in Luria-Bertani (LB), Brain Heart Infusion (BHI), and M9 or M8 minimal media plates solidified with 1% agar. For Kp monoculture, KPPR1 was spread on the entire surface of a 90 mm plate, creating a lawn, while for Pa monoculture, 5 μL of an overnight culture of PA14 was spotted at the center of another 90 mm plate. For coculture experiments, the center of a lawn of Kp was spotted with 5 μL of an overnight culture of Pa (Figure 1A). Kp exhibited profuse growth as a monoculture lawn in 24 hours at 37°C. Pa spot, as monoculture, grew in the center only up to a 4 mm circle under the same conditions. On coculture plates, Pa spot grew to a diameter of ~7 mm while a much larger clearance zone emerged around Pa spot on M9 and M8 plates (Figure 1A). In contrast, no clearance was observed around Pa on LB and BHI coculture plates (Figure 1A). The clearance of Kp from the area surrounding Pa was visualized using GFP expressing Kp and mCherry expressing Pa on M9 plates (Figure 1B).

**Figure 1:**
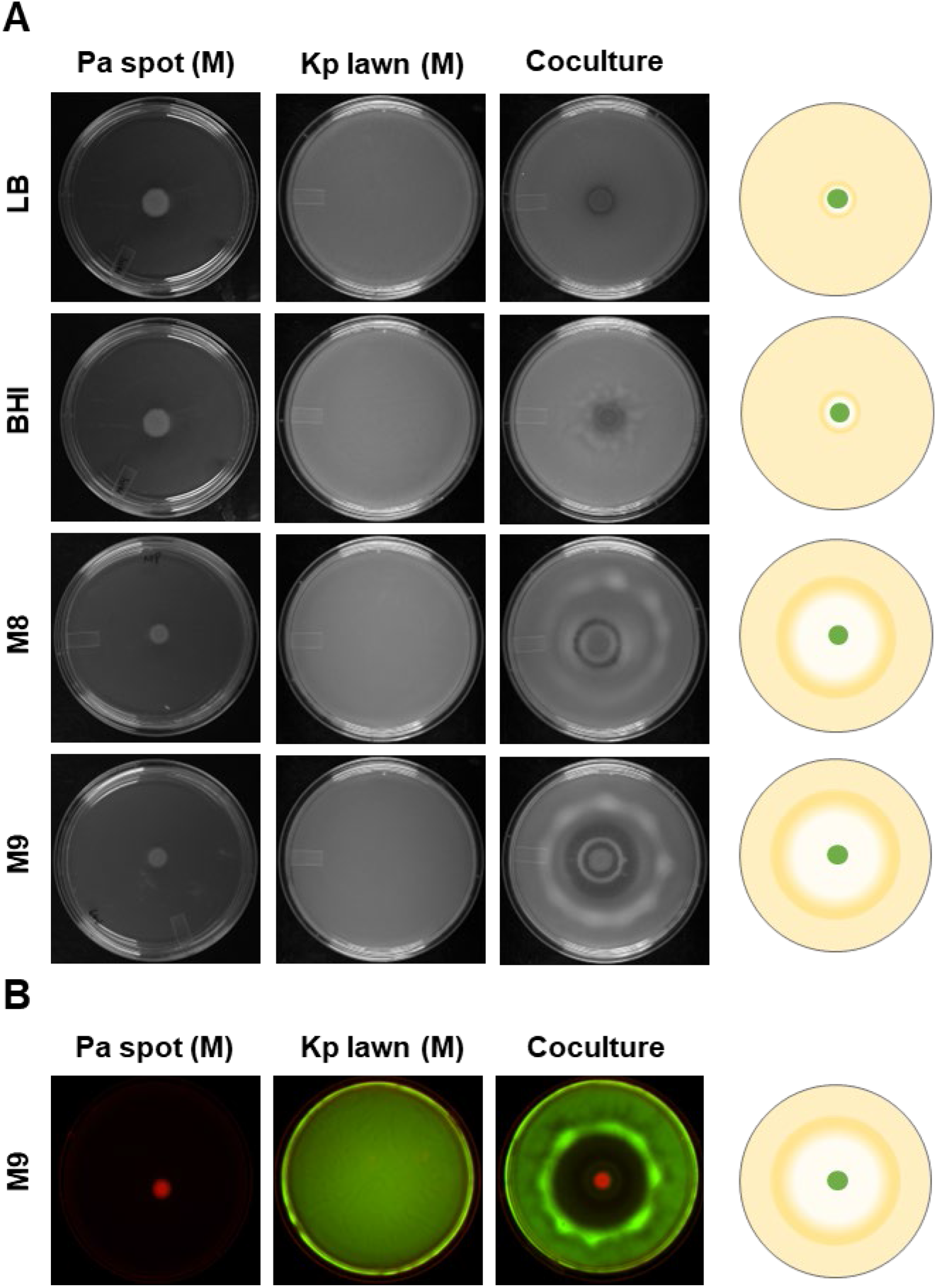
Interaction of *Pseudomonas aeruginosa* and *Klebsiella pneumonia* is media-dependent. (**A)** *P. aeruginosa* (Pa) and *K. pneumoniae* (Kp) Monocultures (M) and Cocultures (C) on LB, BHI, M8, M9 media solidified with 1% agar in 90 mm Petri plate. A schematic representation of the coculture plate is shown on the right side. **(B)** Interaction of *P. aeruginosa* expressing mCherry and *K. pneumoniae* expressing GFP on M9 plate.

To understand whether clearance emerged as a function of Pa or Kp, we also examined the reciprocal interaction wherein we plated Pa cells on the entire agar surface, followed by spotting Kp cells in the center. In this scenario, no zone of clearance appeared (Figure S1A). This indicated that the clearance zone emerged because of the action exerted by *P. aeruginosa*, and Kp does not have specific offensive capability.

To understand whether this behavior is performed explicitly by Pa, we spotted *S. aureus, Escherichia coli* or *Proteus mirabilis* on the lawn of Kp. Still, we did not observe any clearance (Figure S1B), suggesting that the clearance of Kp is a specific function of Pa cells. The minimal media-specific behavior of Pa against Kp is reminiscent of the swarming motility of *P. aeruginosa* which is also greatly influenced by growth media (Badal et al., 2021; Kollaran et al., 2019). Altogether, our experiments indicate that some factor(s) in the M9 medium promotes action of Pa against Kp.

### Toroidal displacement of *K. pneumoniae* by *P. aeruginosa* on minimal media

To better understand the nature of the interaction between Pa and Kp, we closely monitored the development of the clearance zone in coculture plates, in a time-lapse movie (See supplementary movie S1). We observed a wave emanating from Pa cells present in the center (Radial Point or RP) pushing Kp cells outward. This created a clearance zone (CZ) and a mass of material collected in the toroid zone (TZ) (Figure 2B). This indicated that Kp cells were being actively pushed radially outward to form a doughnut or toroid shape, where Kp cells and possibly other materials accumulated, making a ring-like appearance (Figure 2B). We term this active pushing phenomenon to form a doughnut or toroid shape as *‘toroidal displacement’*.

**Figure 2:**
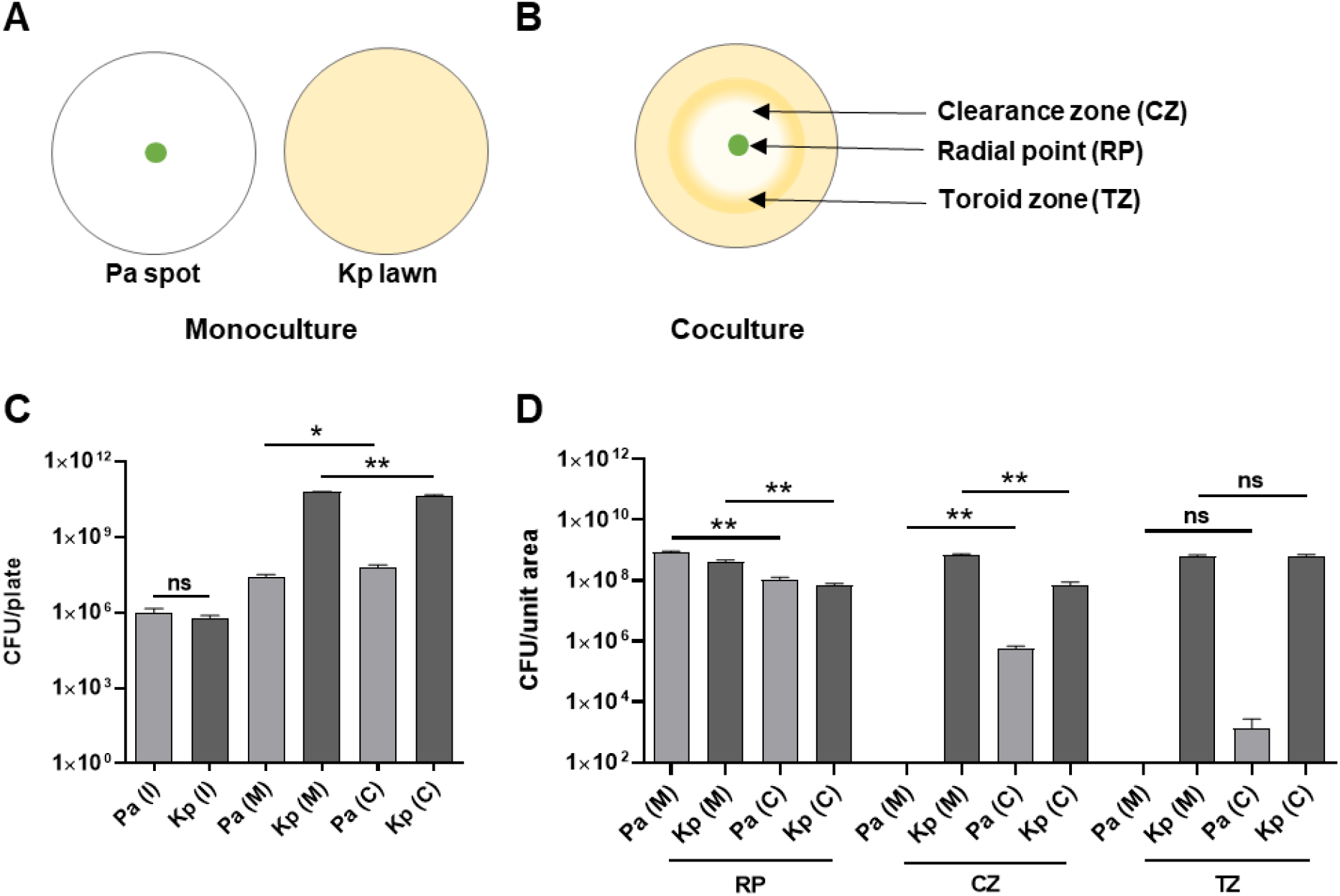
Toroidal displacement of *K. pneumoniae* by *P. aeruginosa* on a solid surface. **(A)** Schematic representation of Pa and Kp monoculture plates. (**B)** Schematic representation of Pa and Kp on M9 coculture plates showing different zones, Radial Point (RP), Clearance Zone (CZ), and Toroid Zone (TZ). **(C)** Enumeration of total CFU of Pa and Kp from inoculum (I), monoculture (M) and coculture (C) plates. **(D)** Enumeration of CFU of Pa and Kp from different zones of monoculture (M) and coculture (C) plates. An unpaired *t*-test was used for the analysis of significance (ns, non significant; *, P≤0.05; **, P≤0.01).

To understand the effect of interactions on the population of these bacteria, we harvested total cells from the entire mono and coculture plates and enumerated colony forming units (CFU) for each bacterium using selective media. We observed a small increase in the Pa population and decrease in Kp population in coculture (Figure 2C). We also harvested cells from a ~38 mm^2^ area of various regions (RP, CZ and TZ) of coculture plates at 24 hours and enumerated CFU for Pa and Kp. We enumerated CFU from corresponding zones of Pa and Kp monoculture plates as respective controls. We observed that Kp population at the radial point and clearance zone were slightly less in coculture plates than in Kp monoculture plates (Figure 2D) consistent with them being pushed out. Interestingly, the Pa population in the clearance zone of coculture plates was more than in the Pa monoculture control plates suggesting that some Pa cells could move into the clearance zone. In the toroid zone (TZ), where there is increased material, we could barely detect Pa cells, but we found plenty of Kp cells. To check if either bacterium was using bactericidal offense mechanisms, we performed a live-dead assay, taking cells from different zones of the coculture plate. Although we observed 100% death in heat-killed bacteria (used as control), we observed a few dead, propidium iodide-stained cells in RP and CZ regions, suggesting negligible death in coculture plates (Figure S2). These results indicated that Pa employs a non-bactericidal mechanism to create the clearance zone.

### Quorum sensing in *P. aeruginosa* facilitates toroidal displacement of *K. pneumoniae*

Since Quorum Sensing (QS) modulates offense and defense strategies of *P. aeruginosa* (Whiteley et al., 1999), we first asked if quorum is necessary for toroidal displacement. We studied the requirements of various components of hierarchical QS systems in *P. aeruginosa* (Lee & Zhang, 2015). We found that RhlR, the transcriptional regulator of the autoinducer butanoyl homoserine lactone (C_4_HSL), as well as RhlI, an enzyme for the autoinducer, were needed in Pa for the optimal displacement of the Kp population (Figure 3A). Reduction in displacement of Kp was also observed upon spotting mCherry expressing *rhlR* mutant on GFP expressing Kp (Figure 3B). The reduced displacement was also reflected in CFU analysis. The total population of Kp between mono and coculture plates were similar while there was small but significant decline in the population of Pa (*rhlR*) cells in coculture plates (Figure 3C). We also examined the requirement for LasR/I system consisting of LasR, transcriptional activator and LasI involved in the synthesis of dodecanoyl homoserine lactone, autoinducer C_12_HSL (Pearson et al., 1997). We observed that both LasI and LasR were dispensable for the clearance of Kp (Figure S3). Neither LasA nor neighboring LasB proteases of Pa, both under the control of the LasR/I system, were required for the clearance of Kp (Figure S3). These results suggest that only RhlR/I quorum sensing system is necessary to drive clearance of Kp in a surface interaction regime.

**Figure 3:**
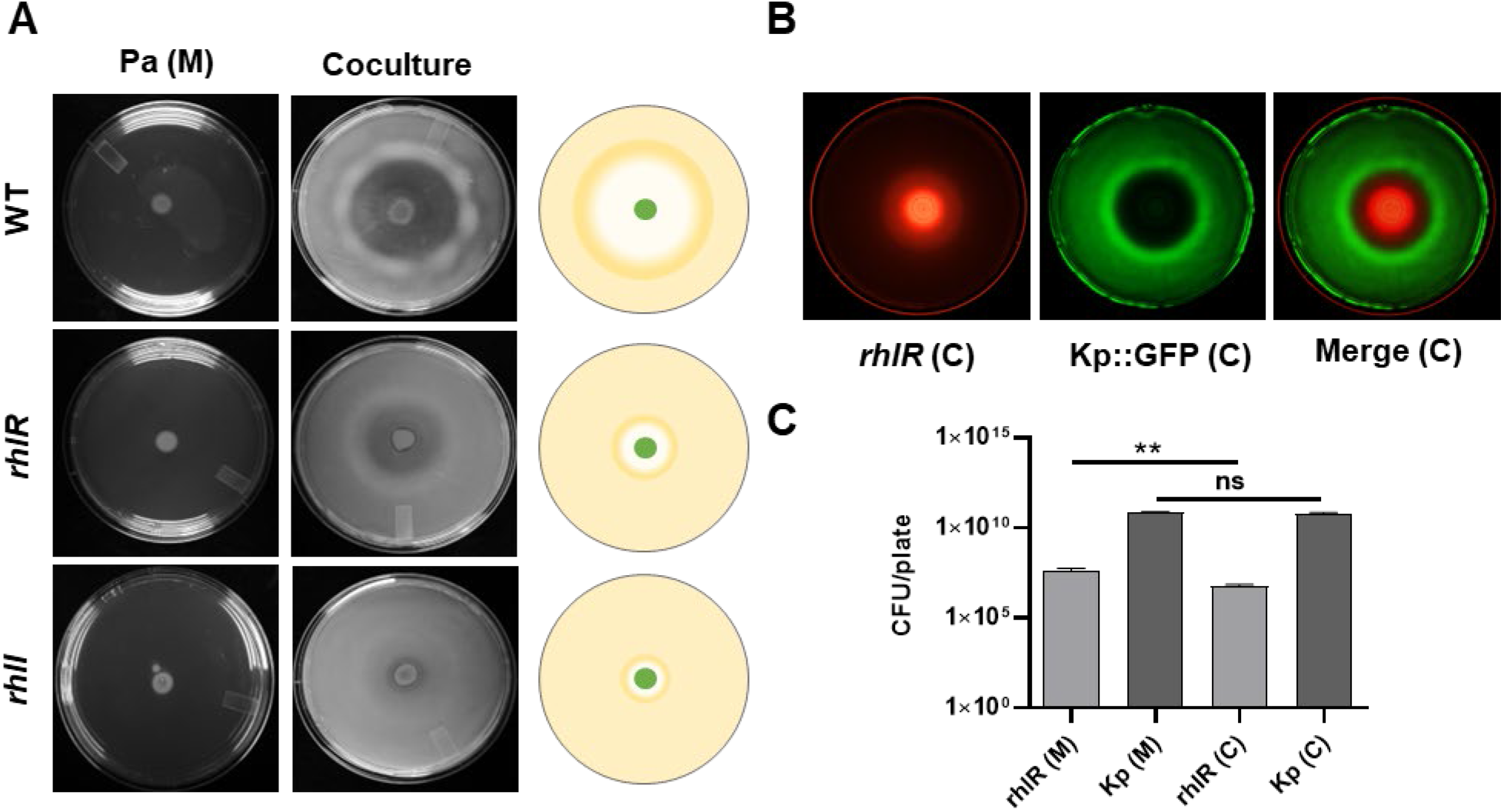
RhlR/I quorum sensing system of *P. aeruginosa* is necessary for displacement of *K. pneumoniae*. **(A).** Interaction of wild type, *rhlR* and *rhlI* mutants of *P. aeruginosa* with Kp on M9 coculture assay plate. Pa (M) represents a Pa monoculture plate. **(B)**. Interaction of *rhlR* expressing mCherry with Kp expressing GFP on M9 coculture plate. **(C)**. Enumeration of total CFU of Pa and Kp from monoculture (M) and coculture (C) plates. An unpaired *t*-test was used for the analysis of significance (ns, non significant; **, P≤0.01).

### Rhamnolipids drive toroidal displacement activity of *P. aeruginosa*

To understand if a specific set of virulence factors of Pa drive the toroidal displacement of Kp cells, we examined virulence factor (Chen et al., 2005) mutants of *P. aeruginosa* in coculture assays. We tested 188 transposon insertion mutants against Kp (Table S2). Surprisingly, we found no evidence for the involvement of most virulence factors, including phenazines, flagella, lipopolysaccharide, pili, toxins, alginate etc. This analysis indicated that Pa does not utilize bactericidal mechanisms for clearance of Kp, consistent with our live/dead analysis in coculture plates. Pa likely utilizes killing-independent mechanisms to push Kp cells out of the way.

Of the 188 factors tested, only two were necessary for the optimal displacement of Kp. We found that mutations in either *rhlA* or *rhlB* genes, encoding rhamnosyl transferase I necessary for synthesizing rhamnolipid biosurfactant, suppressed the toroidal displacement of Kp (Figure 4A). This indicated that rhamnolipids help Pa push Kp to the edge of the clearance zone. To confirm that Kp is indeed pushed out by Pa, we studied whether small spots of Kp::GFP over the lawn of unlabeled Kp (Figure 4B) disappear or get displaced after 24 hours. We found that fluorescent Kp spot was pushed out all the way to the toroid zone but there was no decline in fluorescence suggesting there was no death of Kp. *rhlA* and *rhlB* mutants of Pa were unable to push Kp::GFP (Figure 4B). These experiments provided conclusive evidence that Pa actively pushes Kp and it does so in a surfactant-dependent manner.

**Figure 4:**
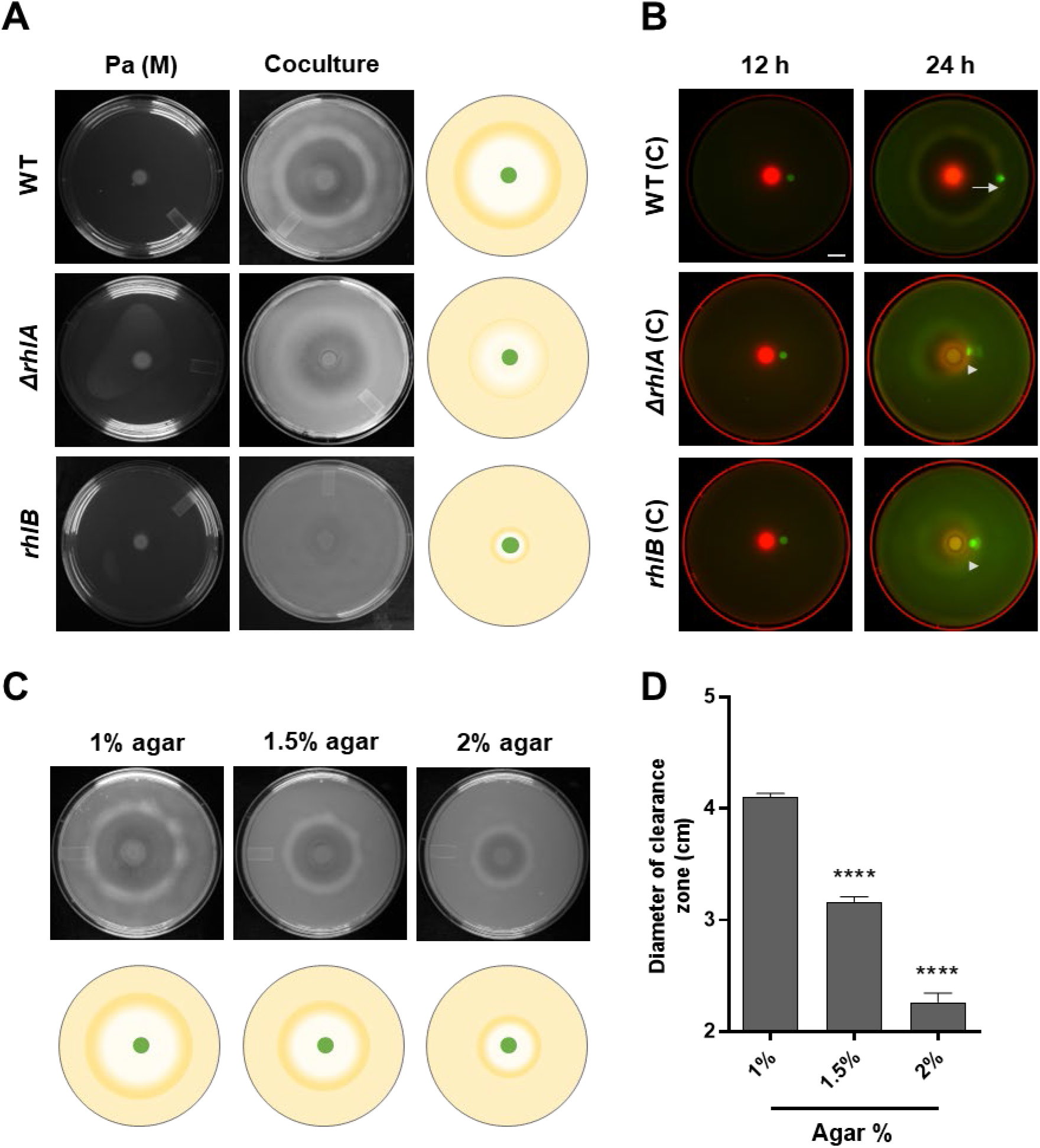
*P. aeruginosa* employs rhamnolipids for the toroidal displacement of *K. pneumoniae*. **(A)**. Interaction of Pa (wild type, *rhlA* and *rhlB*) with Kp on M9 coculture plates. Pa (M) represents a Pa monoculture plate. **(B)**. Location of GFP expressing Kp, spotted 1 cm from the center of a Pa-untagged Kp coculture plate, at 12 hours and 24 hours of incubation. Displacement of Kp::GFP spot is indicated with an arrow in 24 hour images. Scale bar, 1 cm. **(C)**. Pa-Kp coculture assays on M9 plate solidified with 1, 1.5 and 2% agar. Schematic representations for the same is shown. **(D)**. Diameter of the clearance zone on Pa-Kp coculture assays on M9 plate solidified with 1, 1.5 and 2% agar. An unpaired *t*-test was used for the analysis of significance (****, P ≤ 0.0001).

To test whether the availability of water also determines the pushing ability of Pa, we performed coculture experiments on plates with increasing concentration of agar. As shown in Figures 4C and 4D, there was significant decrease in the diameter of the clearance zone for Kp with an increase in agar percentage from 1% to 2% suggesting that availability of water positively affects pushing ability of Pa. Taken together, our analysis suggested that the displacement of Kp by Pa is mediated by rhamnolipid biosurfactant.

### Iron limitation promotes toroidal displacement action of *P. aeruginosa* against *K. pneumoniae*

Iron is required for respiration and growth in all living organisms, and iron limitation is one of the significant drivers of virulence in *P. aeruginosa* (Meyer et al., 1996; Minandri et al., 2016; Vasil & Ochsner, 1999). Therefore, we asked whether iron limitation drives toroidal displacement of Kp by Pa. We examined the toroidal displacement activity on coculture plates supplemented with an increasing concentration of iron (ferrous sulphate, Fe_2_SO_4_. 7H_2_O) ranging from 2 μM to 10 μM in addition to un-supplemented control. We found that iron above 2 μM suppressed toroidal displacement of Kp while 10 μM iron was sufficient to completely abolish toroidal displacement (Figure 5A). To confirm the role of iron limitation, we added an iron chelator, 2,2’-Bipyridine (BP), to iron-supplemented coculture plates. We could restore toroidal displacement of Kp by adding 125 μM BP in iron-supplemented plates (Figure 5B).

**Figure 5:**
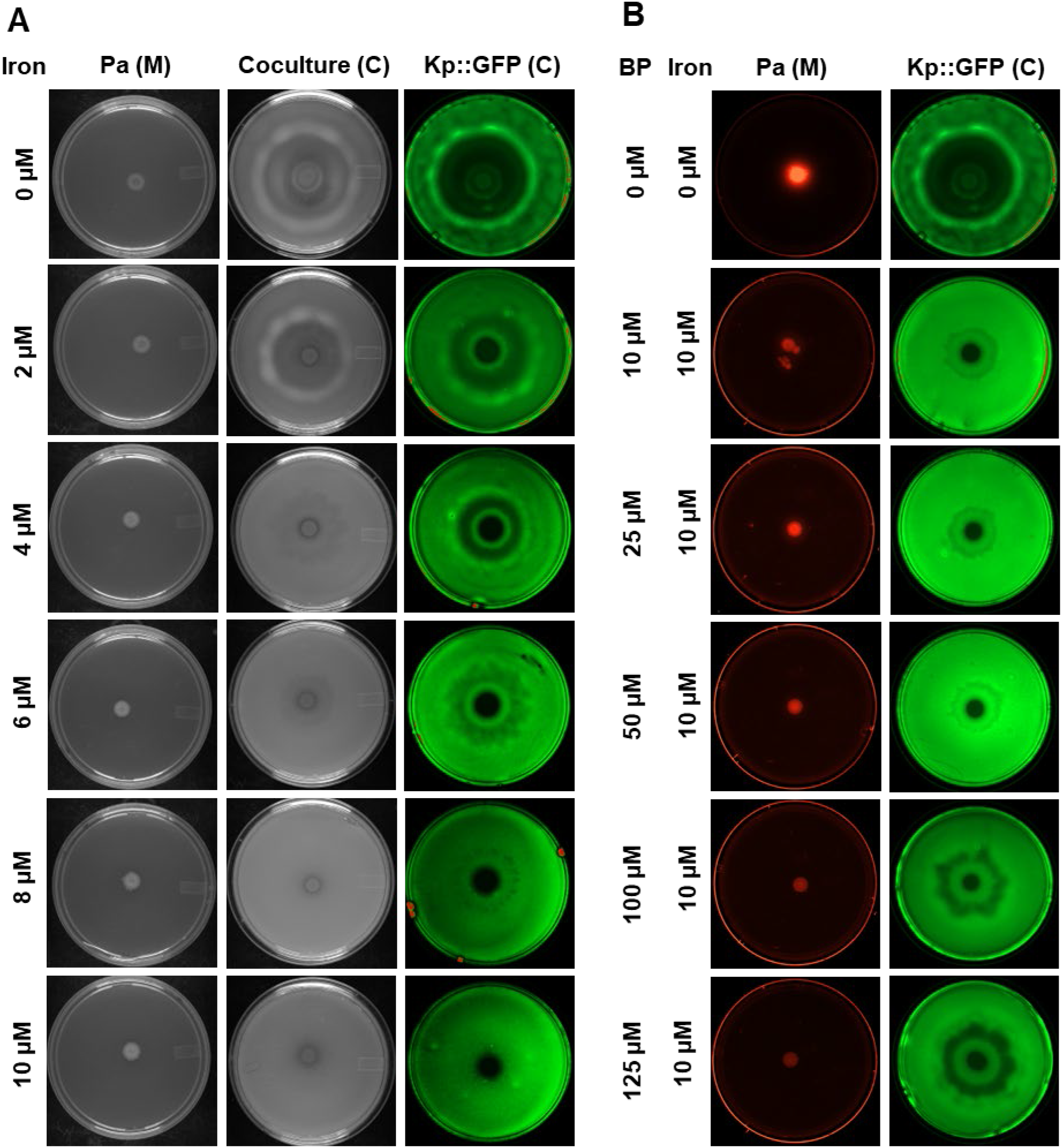
Iron-limitation promotes toroidal displacement of *K. pneumoniae* by *P. aeruginosa*. **(A)** Pa-Kp coculture assays on M9 plates supplemented with 0 to 10 μM of FeSO_4_.7H_2_O. **(B)** Pa-Kp coculture assays on M9 plates supplemented with 10 μM of FeSO_4_.7H_2_O, and 0 to 125 μM of Iron chelator 2,2’-Bipyridyl (BP). Pa (M) represents Pa monoculture, and Kp::GFP (C) represents Pa-Kp::GFP coculture plate.

The analyses of toroidal displacement with additional iron and iron chelator provided strong evidence that iron limitation drives the toroidal displacement of *K. pneumoniae* by *P. aeruginosa*.

### Competition for iron sets off an offense response in *P. aeruginosa* against *K. pneumoniae*

All living organisms, including bacteria, employ various strategies to harvest iron from the environment and their neighbors. Siderophores are small organic molecules secreted by bacteria to harvest extracellular iron (Judith & Manuela, 2016). Pyoverdine is one of the major siderophores secreted by Pseudomonas. We examined the expression of various genes involved in the synthesis of pyoverdine in *P. aeruginosa* schematic in Figure S4A). We examined the level of transcripts for *pvdG, pvdA, pvdF, pvdE, pvdQ, pvdP, pvdR, fpvA, fpvE, fpvK, pvdS* in *P. aeruginosa* grown in LB or M9 medium. We found several fold upregulations in all these genes in cells grown in M9 compared to cells grown in LB medium (Figure 6A). This indicated that the M9 medium has less iron than the LB medium. Independent measurement of pyoverdine secreted by Pa in M9 and LB confirmed that M9 media promotes pyoverdine production (Figure S5). *pvdE* mutant, defective in the export of pyoverdine, had very little secreted pyoverdine as expected (Figure S5). When we supplemented M9 medium with 36 μM iron, we observed suppression in the expression levels of transcripts for genes involved in siderophore synthesis in Pa (Figure 6C).

**Figure 6:**
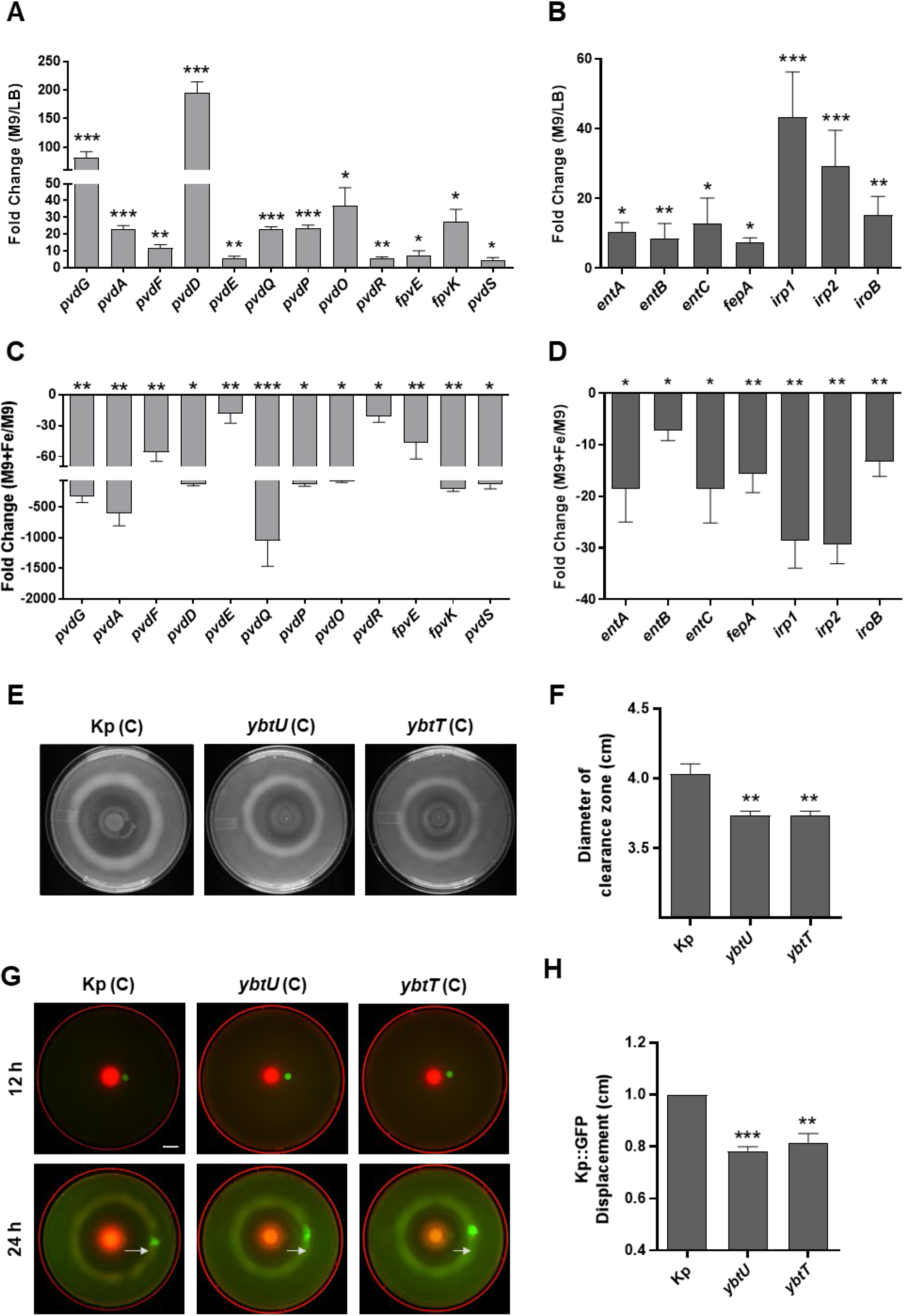
*P. aeruginosa* and *K. pneumoniae* compete for iron in M9 medium. Analysis of expression of genes involved in siderophore synthesis and export in *P. aeruginosa* **(A)** grown in M9 medium over LB medium, or **(C)** grown in iron supplemented M9 medium over M9 medium for 24 hours at 37°C. Analysis of expression of genes involved in siderophore synthesis in *K. pneumoniae* **(B)** grown in M9 medium over LB medium, or **(D)** grown in iron supplemented M9 medium over M9 medium for 24 hours at 37°C. **(E)** Interaction of Pa with Kp (wild type, *ybtU* and *ybtT*) on M9 coculture plates and **(F)** Diameter of the clearance zones in respective assays. **(G)** Location of GFP expressing Kp, spotted 1 cm from the center of a Pa-Kp, Pa-*ybtU* or Pa-*ybtT* coculture plate, at 12 hours and 24 hours of incubation at 37°C. Scale bar, 1 cm. **(H)** Displacement of spot of GFP expressing Kp on Pa-Kp, *Pa-ybtU* or Pa-*ybtT* coculture plate after 24 hours. An unpaired *t*-test was used for the analysis of significance (*, P ≤ 0.05; **, P ≤ 0.01;***, P≤ 0.001).

*K. pneumoniae* is known to synthesize as many as 4 different siderophores namely enterobactin, yersiniabactin, salmochelin and aerobactin to establish infection (Paczosa & Mecsas, 2016) (schematic in Figure S4B). We examined the expression of some of the genes involved in the synthesis of enterobactin (*entA, entB, entC*, and *fepA*), Salmochelin (*iroB*), and yersiniabactin (*irp1, irp2*) and we found that all the transcripts were upregulated in Kp grown in M9 medium over Kp grown in LB medium (Figure 6B). This upregulation was suppressed when Kp cells were grown in M9 medium supplemented with 36 μM of iron (Figure 6D). These experiments suggested that both Pa and Kp respond to iron-limiting conditions in the M9 medium by upregulating genes involved in the production of various siderophores.

To understand the importance of iron in driving interbacterial interactions, we measured total iron available in M9 and LB medium using an iron estimation kit. While iron was undetectable in the M9 medium, there was up to 2 μM of free iron in the LB medium. To get an insight into the ability of Pa and Kp to utilize iron, we grew Kp and Pa individually in the LB medium and measured iron in the spent media after 6, 12 and 16 hours of growth. As shown in Figure S6, both bacteria depleted iron, in a time-dependent manner. To reduce the competition for iron, we studied *ybtT* and *ybtU* mutant of Kp (lacking yersiniabactin siderophore). As expected, the clearance zone for *ybtU* and *ybtT* was smaller than the clearance of parental KPPR1 in coculture plates (Figure 6, E-F). We also quantified displacement of Kp::GFP by Pa, and found that displacement was lower on *ybtU* and *ybtU* mutants of Kp (Figure 6, G-H).

Altogether, our experiments indicate that iron competition alters the behavior of *P. aeruginosa* in a manner that allows it to push *K. pneumoniae* out of its way.

### Rhamnolipid production induced by iron limitation regulates *P. aeruginosa* behavior on solid surfaces

Our study suggested that iron limitation is the driving force for the offense response in Pa against Kp. We also observed that Pa employs a rhamnolipid-dependent mechanism to push Kp to the toroid zone. To establish a cause-and-effect link between these two events, we asked if iron limitation drives rhamnolipid synthesis. We analyzed the expression of transcripts for rhamnosyl transferases, *rhlA* and *rhlB*, in Pa cells grown in LB and M9. We found that cells grown in the M9 medium had 3-4 fold increased expression of *rhlA* and *rhlB* than cells grown in the LB medium (Figure 7A). Further, we observed that iron supplementation in M9 medium completely suppressed the upregulation of *rhlA* and *rhlB* (Figure 7B). We further confirmed the link between iron limitation and rhamnolipids in an orthogonal assay. Since rhamnolipid is essential for swarming in Pa (Caiazza et al., 2005), we predicted that iron supplementation would suppress rhamnolipid production and thus will suppress swarming. As shown in Figure S7, Pa swarms on M9 medium solidified with 0.6% agar as shown earlier (Badal et al., 2021; Kollaran et al., 2019). However, supplementation of M9 swarm agar plates with iron suppressed swarming in a dose-dependent manner (Figure S7).

**Figure 7:**
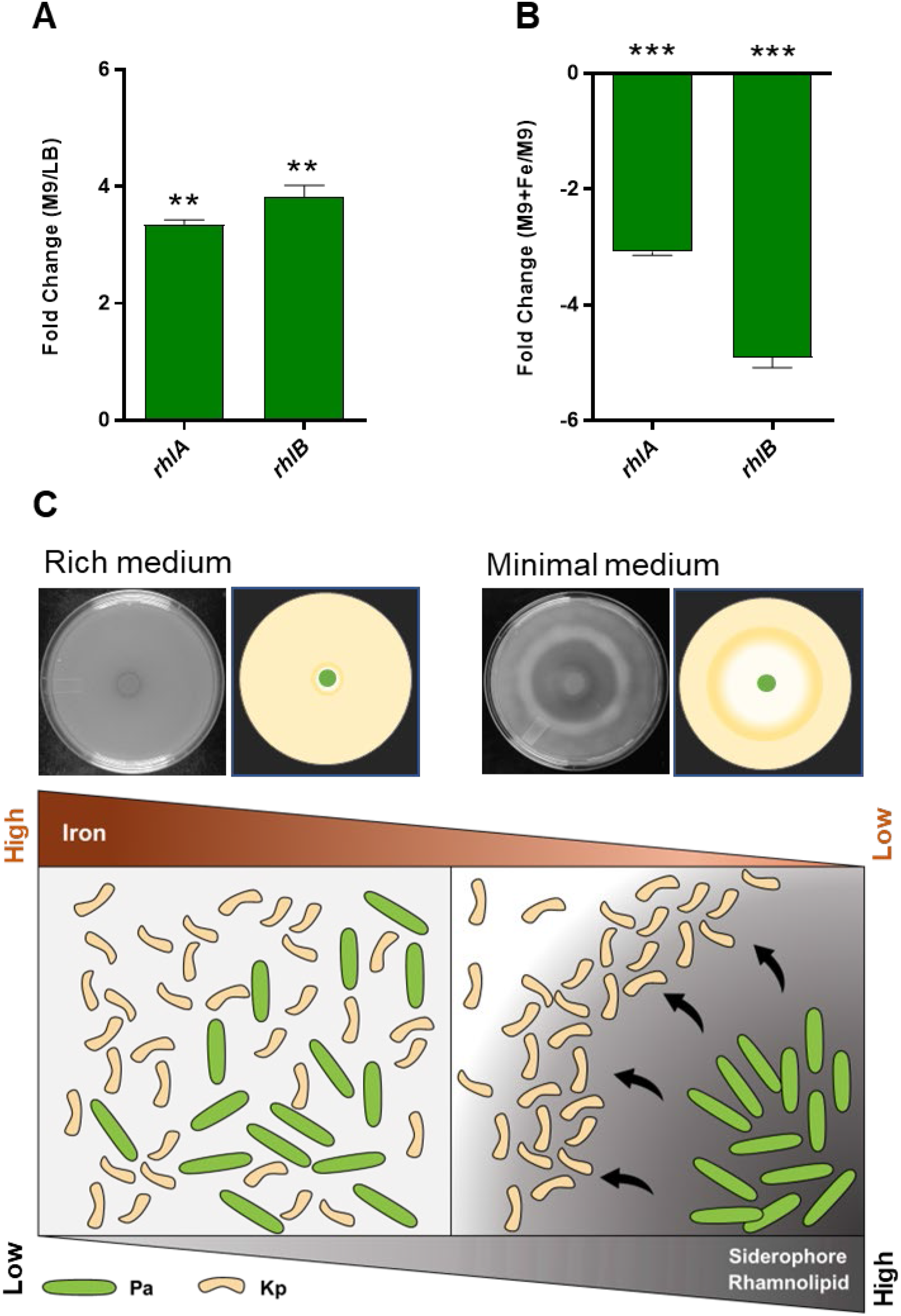
Rhamnolipid production induced by iron limitation regulates *P. aeruginosa* behavior on surfaces. **(A)** Analysis of expression of *rhlA and rhlB* genes of *P. aeruginosa* after 24 hours of growth in M9 medium over LB medium, and **(B)** in iron-supplemented M9 medium over M9 medium alone. An unpaired t-test was used for the analysis of significance (**, P ≤ 0.01; ***, P ≤ 0.001). **(C)** A model for the interaction between *P. aeruginosa* and *K. pneumoniae*. These bacteria can co-exist in nutrient-replete conditions. Under iron-limiting conditions, siderophore synthesis is induced in both bacteria. Iron limitation can also turn on the synthesis of biosurfactants, rhamnolipids, in *P. aeruginosa* allowing it to push away *K. pneumoniae*.

Altogether, our study provides strong evidence that iron limitation promotes rhamnolipid production in *P. aeruginosa*, allowing it to push *K. pneumoniae* out of its way, to limit iron competition. A model for the interaction of *P. aeruginosa* and *K. pneumoniae* under iron-limiting conditions is shown in Figure 7C.

## DISCUSSION

Studies on the interactions of *P. aeruginosa* with its neighbors have shown that it employs a range of offense and defense mechanisms including use of soluble toxic molecules such as phenazines, hydrogen cyanide, proteases, phospholipases against Gram positive bacteria, and pathogenic yeast. In this study, we have shown that under specific conditions, *P. aeruginosa* does not utilize bactericidal mechanisms. Rather, it adopts a unique pushing mechanism to displace *K. pneumoniae*, a fierce scavenger of iron. Careful analyses of the conditions suggest that *P. aeruginosa* can co-exist with *K. pneumoniae* in rich media but under the condition of iron limitation, it pushes *K. pneumoniae* out using rhamnolipid biosurfactant.

Our study found no evidence that *P. aeruginosa* deploys bactericidal mechanisms against *K. pneumoniae*. Why should this be so? One reason could be the innocuous nature of *K. pneumoniae* against *P. aeruginosa*. Kp does not attempt to kill or get rid of Pa (Figure S2) and thus draws no defense response from Pa. A second reason could lie in the features of *K. pneumoniae* cells. Perhaps the thick layer of carbohydrate-rich capsule surrounding cells (Mike et al., 2021) renders them impervious to soluble toxic molecules of *P. aeruginosa* making them ineffective. In such a scenario, synthesis and secretion of virulence factors by *P. aeruginosa* would be a futile and expensive endeavor.

Why does iron limitation drive toroidal displacement response from *P. aeruginosa?* While most organisms have ways to harvest iron from their surroundings, *P. aeruginosa* and *K. pneumoniae* excel at iron scavenging. In response to low Fe^2+^, Pa produces a high-affinity siderophore called pyoverdine allowing it to scavenge Fe^3+^ from the environment and promotes its own growth (Minandri et al., 2016; Takase et al., 2000). In addition, *P. aeruginosa* can utilize siderophores from other bacteria in a process called siderophore piracy. Most Klebsiella species also have an impressive repertoire of at least three siderophores, enterobactin, salmochelin and yersiniabactin (Holden et al., 2018). These two bacteria rapidly utilize iron in their surrounding (Supplementary Figure S6) setting off competition for iron in minimal media. Depletion of iron not only drives siderophore synthesis in *P. aeruginosa* (Figure 6A, 6C), it also induces the synthesis of rhamnolipids (Figure 7, A-B). Thus, the biosurfactant circuit has evolved to be responsive to iron availability allowing *P. aeruginosa* to use the surfactant for driving competitors for iron away.

While *P. aeruginosa* and *K. pneumoniae* are two of the fierce scavengers of iron, the same is not true for other microbes such as *S. aureus, E. coli* or *C. neoformans* (Hammer & Skaar, 2011; Hartmann & Braun, 1981; Vartivarian et al., 1995). We found very little iron depletion by *S. aureus* (data not shown). The low iron scavenging capacity of other microbes (*S. aureus* or yeast) probably explains the lack of toroidal displacement in their interaction with *P. aeruginosa* (Badal et al., 2021; Hotterbeekx et al., 2017; Mashburn et al., 2005). If *P. aeruginosa* does not experience iron limitation, there will not be enough surfactants to cause toroidal displacement. This is consistent with our experiments with siderophore-deficient *K. pneumoniae* strains. Both *ybtU* or *ybtT* strains of *K. pneumoniae* experienced delayed toroidal displacement response from *P. aeruginosa*.

Microbial biosurfactants are expensive to make but facilitate major processes such as (i) the dispersal of bacteria from biofilms allowing them to colonize new niches, and (ii) reduction of surface tension for swarming motility. Our study suggests that rhamnolipids can also aid in the competitive fitness of *P. aeruginosa* by reducing competition for nutrients such as iron. *P. aeruginosa* likely utilizes its surfactant to reduce adhesion between *K. pneumoniae* cells with the agar substratum. It is possible that *P. aeruginosa* can utilize surfactants to perform toroidal displacement in response to competition for other nutrients besides iron. A careful analysis of *P. aeruginosa* behaviors under various physiologically relevant, nutrient-limiting conditions will elucidate environmental features which impinge on biosurfactant production in bacteria.

## METHODS

### Microbes and growth conditions

*Pseudomonas aeruginosa* PA14 (WT) and the isogenic transposon-insertion mutant strains (Liberati et al., 2006) were obtained from Prof. Frederick Ausubel. Additional strains used in this study are listed in tables S1 and S2. Unless otherwise mentioned, PA14 was routinely cultured in LB broth at 37°C, while the mutants were cultured in LB broth with 50 μg mL ^-1^ gentamicin. Similarly, *E. coli* strains used for cloning experiments were routinely cultured in LB broth at 37°C with appropriate antibiotics. The *Klebsiella pneumoniae* KPPR1 strain and transposon insertion mutants, *ybtU* and *ybtT*, were obtained from Prof. Harry Mobley (Mike et al., 2021). KPPR1 was cultured in LB broth at 37°C, while the transposon-insertion mutants were cultured in LB broth with 50 μg mL ^-1^ kanamycin. Kp::GFP strain were obtained from Prof. Gad Frankel (Wong et al., 2019). It was cultured in LB with 50 μg mL ^-1^ carbenicillin at 37°C. *S. aureus*, *E. coli* or *P. mirabilis* were cultured in LB at 37°C. Additional media used in this study include BHI (HiMedia), M9 (8.6 mM NaCl, 20 mM NH_4_Cl, 1 mM CaCl_2_,1 mM MgSO_4_, 22 mM KH_2_PO_4_, 12 mM Na_2_HPO_4_, 0.2% glucose and 0.5% casamino acids). M8 is a modification of the M9 medium (excluding NH_4_Cl and CaCl_2_ salts) (Kohler et al., 2000). Fe_2_SO_4_.7H_2_O was used as the source of iron in all the experiments involving iron. Iron chelator, 2,2’ Bipyridine (Sigma-Aldrich) was used at indicated concentrations.

### Surface Interaction assay

Kp lawn was made using 1:100 dilution of a 0.5 OD_600_ culture of Kp. 500 μL of the diluted culture was spread on LB, BHI, M9 or M8 medium plates solidified with 1% Bacto agar (BD) and excess liquid was removed with a pipette. The lawn was allowed to dry for 40 minutes at room temperature. 5 μL of 2.0 OD_600_ culture of Pa was spotted at the center of Kp lawn (coculture) or a plain plate (monoculture). Plates were dried for 20 more minutes and then incubated at 37°C for interaction to take place. Both mono and coculture plates were imaged in Vilber E-box after 24 hours of incubation. Images involving fluorescence were captured in the Bio-Rad ChemiDoc MP system.

### Colony Forming Unit (CFU) assay

CFU was estimated in coculture or monoculture plates by washing off the plate with Phosphate-Buffered Saline (PBS). Serial dilutions were made and plated on cetrimide agar (HiMedia) and HiCrome™ Klebsiella selective Agar Base (HiMedia) to enumerate Pa and Kp respectively. To estimate the numbers of Pa and Kp in different zones (radial point, clearance zone, toroid zone) (Figure 2B) in coculture, a 38 mm^2^ plug of agar was captured using the back of a 1 mL tip. The bacteria from the plug were diluted using PBS and plated on selective agar to enumerate colonies of Pa and Kp.

### Live-dead staining assay

Bacterial cells were collected from different zones of mono and coculture plates after 24 hours of incubation. Live/dead Double Staining Kit (Merck Life Sciences) was used to stain cells followed by microscopy. Heat-killed Kp cells are used as a control. After staining, cells were imaged using a Stedycon optical setup (Abberior Instruments GmbH) integrated with Leica SP 5 microscope (Leica Microsystems GmbH, Manheim, Germany).

### Pyoverdine estimation

A 1.0 OD_600_ Pa or Δ*pvdE* culture was inoculated in M9 and LB broth at 1% inoculum. After growth for 24 hours at 37°C with shaking (180 rpm), cell free supernatants were harvested. Pyoverdine fluorescence was measured in the supernatants using TECAN Infinite M200PRO microplate reader (Zhang & Rainey, 2013)(Ex/Em; 365/460 nm).

### Iron estimation

For iron estimation, 1% inoculum of 1.0 OD_600_ Pa and Kp culture was inoculated in LB broth. Total iron was measured in the spent media after 6, 12 and 16 hours of bacterium growth for 24 hours at 37°C with shaking (180 rpm) using the Iron Assay Kit (Sigma-Aldrich). TECAN infinite M200PRO microplate reader was used for measuring Colorimetric readings.

### Swarming motility assay

Swarming motility assays were performed as described earlier (Kollaran et al., 2019). M9 medium was solidified with 0.6% Bacto agar (BD) and stored for 16 hours at room temperature. All plates were inoculated at the center with 2 μL of overnight Pa culture in LB broth (OD_600_ = 2.8 - 3.0) and incubated at 37°C for 24 hours. Images of the swarm were captured using Vilber E-box.

### RNA preparation

Total RNA was extracted from a final volume of 2 mL of harvested cells from an overnight bacterial culture grown in M9 and M9 supplemented with Fe (normalized to OD_600_ of 1) by the hot-phenol method as described (Varshney et al., 1988). Briefly, cells chilled on the ice were centrifuged at 5000 rpm for 5 minutes. The pellet was resuspended in the lysis buffer (20 mM Tris.HCl (pH 7.5), 20 mM NaCl, 5 mM Na2EDTA, 5 mM VRC, SDS (1% w/v) and 2-Mercaptoethanol (0.7% v/v)) to which equal volume of hot phenol (pH 4.3, 65°C) was added and vortexed, followed by incubation at 65°C for 6 minutes. Samples were centrifuged at 12,000 rpm for 15 minutes to recover the aqueous layer, which was further extracted using an equal volume of phenol:chloroform mixture (12,000 rpm, 10 minutes) followed by an equal volume of chloroform (12,000 rpm, 7 minutes). The RNA was precipitated with ethanol and resuspended in 0.5 mL of molecular-grade water. Subsequently, RNA purity was analyzed spectrophotometrically (NanoDrop™ 1000) and subjected to DNase I treatment.

### qRT-PCR analysis

RNA was treated with DNase I and quantified on a NanoDrop™ spectrophotometer (ND-1000). cDNA was prepared using iScript™ cDNA Synthesis Kit (Bio-Rad). The iTaq Universal SYBR Green Supermix (Bio-Rad) was used or qRT-PCR on Quant Studio 3 Real-time PCR system (Applied Biosystems). We used three biological replicates for each sample and two technical replicates for each biological replicate. Fold change for each transcript was calculated from the C_T_ values normalized to the housekeeping gene *rpoD* following the 2^-ΔΔCT^ method (Livak & Schmittgen, 2001). Primers used for qRT-PCR are described in Supplementary Table S3.

### Construction of Δ*pvdE* strain in *P. aeruginosa* PA14

A marker less deletion strain of *pvdE* was generated by the two-step allelic exchange strategy (Hmelo et al., 2015). Clones were screened by deletion-specific PCR and confirmed by sequencing. See table S3 for primers used for the construction of pEXG2-Δ*pvdE*.

## Data Availability

All data is present in the manuscript and supplementary information.

## Acknowledgement

We thank Gad Frankel, Frederick M Ausubel, Arne Rietsch, Peter Greenberg, Zemer Gitai, and Harry Mobley for reagents and strains. We thank Samay Pande for suggestions on this work. This work was supported by Senior Fellowship from the Wellcome Trust/DBT India Alliance to Varsha Singh (IA/S/21/1/505655). AT is supported by Inspire Fellowship from the Department of Science and Technology, DP is supported by SERB-National Post-Doctoral Fellowship and SP is supported by Kothari Postdoctoral fellowship from UGC, Govt. of India. We thank Prof. Jaydeep K. Basu for facilitating access to the microscope facility at the Department of physics supported by the Ministry of Human Resources Development (MHRD)-funded Institute of Eminence (IOE) project.

## Author Contribution

Conceived and designed the experiments: DP, AT, VS. Performed the experiments: DP, AT. Assisted in microscopy: SP. Analyzed the data: DP, AT, VS. Funding acquisition: VS. Wrote the paper: DP, AT, VS.

