## Supplemental figures for "Toroidal displacement of *Klebsiella pneumoniae* by *Pseudomonas aeruginosa* is a unique mechanism to avoid competition for iron"

Supplementary figures S1 to S7

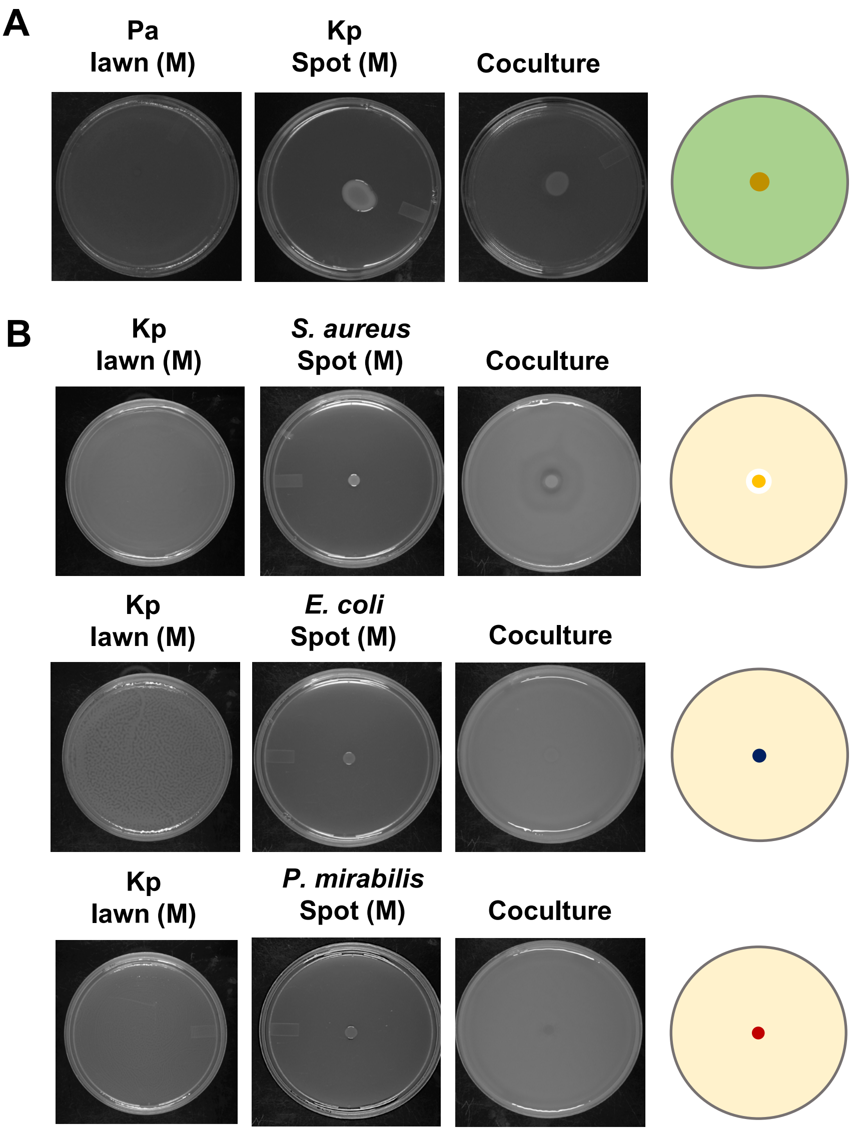


**Figure S1: Toroidal zone formation is due to the activity of *P. aeruginosa*. (A).** Kp spot over Pa lawn in coculture plate. Pa lawn and Kp spot on plain agar plate is used as monoculture (M) control. **(B).** Three different bacteria were spotted on Kp lawn, *S. aureus*, *E. coli*, and *P. mirabilis*. All three bacteria were spotted on plain agar plate for monoculture control.

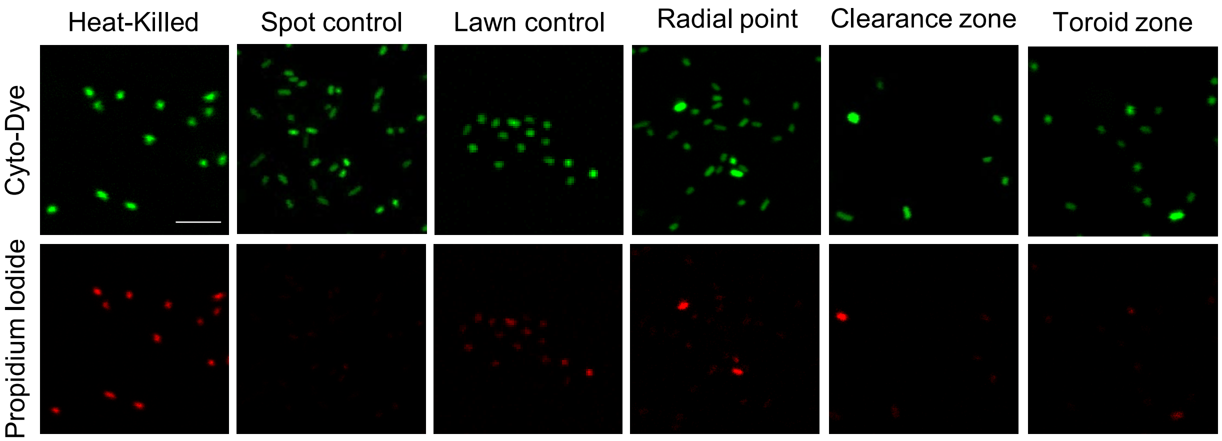


**Figure S2: Live-dead assay shows no sign of killing of *K. pneumoniae* by *P. aeruginosa*.** Live dead cell population was looked in the bacteria population taken from different zones. Heat killed (Positive control), spot control (Pa population), lawn control (Kp lawn population), radial point (bacteria from centre of coculture plate), Clearance zone (bacteria taken from clearance zone), toroid zone (bacteria taken from toroid zone). Scale bar, 2 μm.

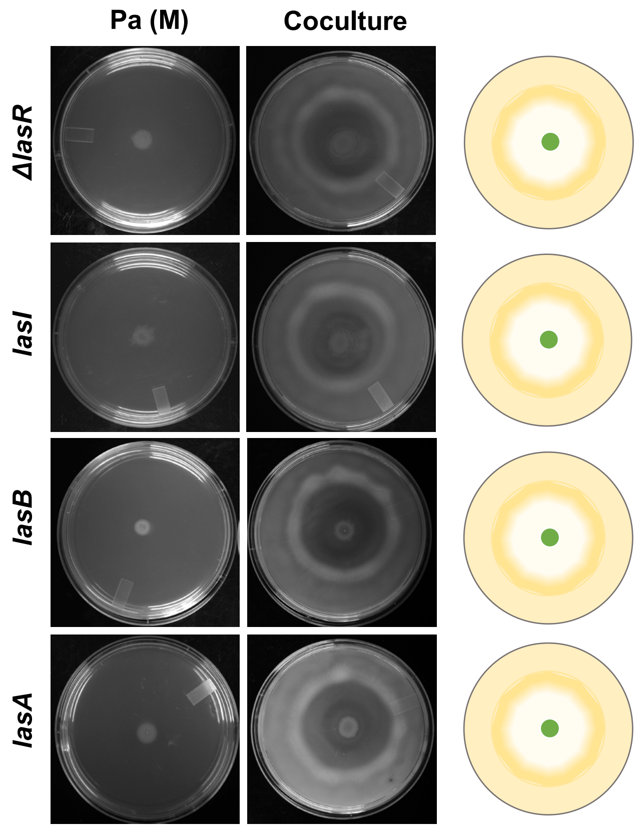


**Figure S3: LasR/I quorum sensing system and its effector toxins are not required for *K. pneumonie* displacement.** Interaction of lasR/I quorum sensing mutants *ΔlasA, lasI, lasB, lasA* with Kp in M9 coculture assay plate.

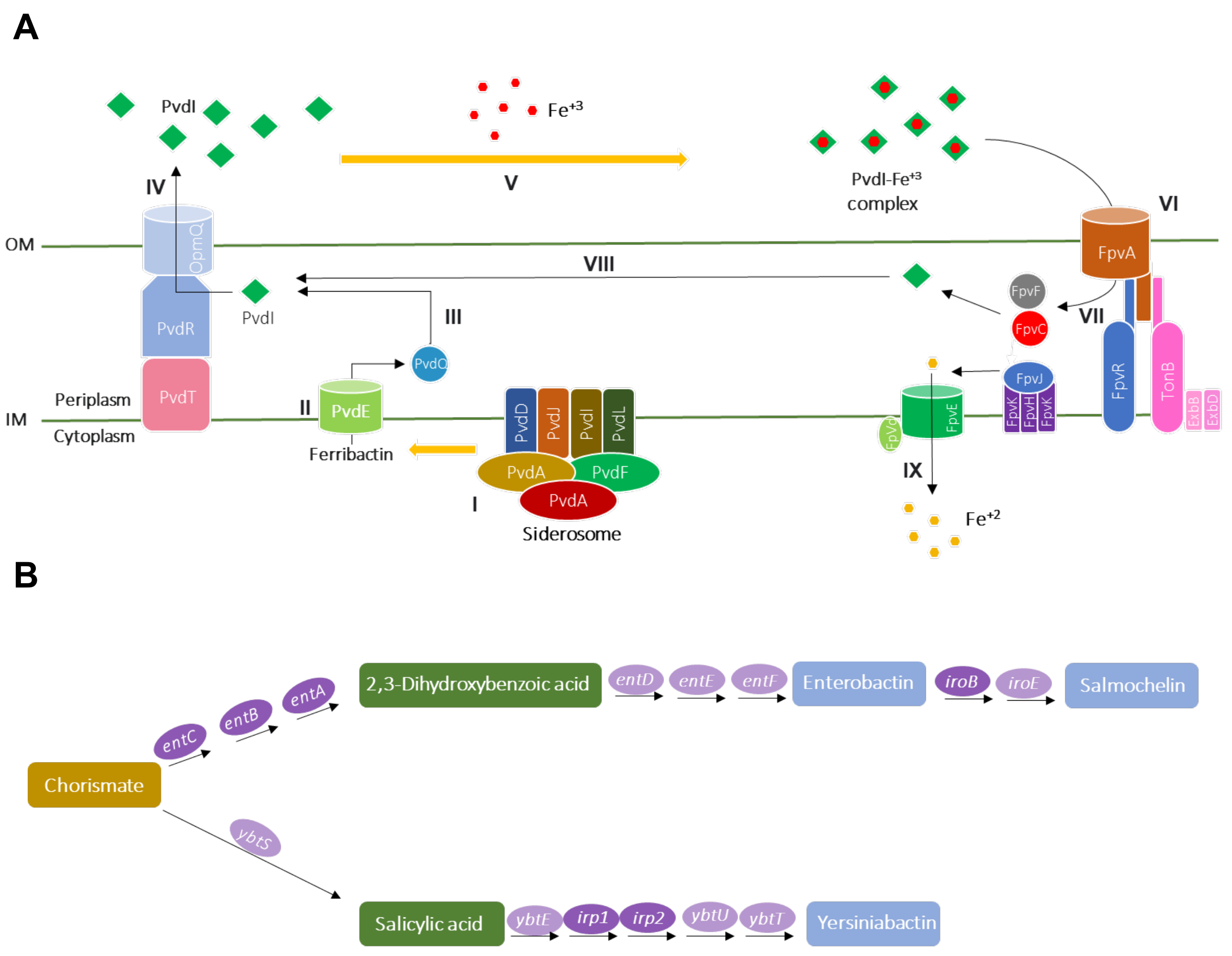


**Figure S4: Schematic representation of *P. aeruginosa* and *K. peumoniae* siderophore synthesis pathways. (A).** Pyoverdine synthesis pathway in Pa. *pvdS* is a sigma factor who to regulates pyoverdine biosynthesis genes. A overview of pyoverdine metabolism is following- I. Synthesis of Pyoverdine precursor (Ferribactin) in siderophore, II. Export into periplasm, III. Pyoverdine maturation, IV. PVDI secretion, V. Iron chelation, VI. Uptake of PvdI-Fe+3, VII. Dissociation of PVDI and iron (Iron reduction), VIII. Recycle, IX. Fe+2 import into cytoplasm **(B).** Kp siderophores synthesis pathway. All three siderophore produced by KPPR1 strain are synthesized from Chorismate (Shikimate pathway).

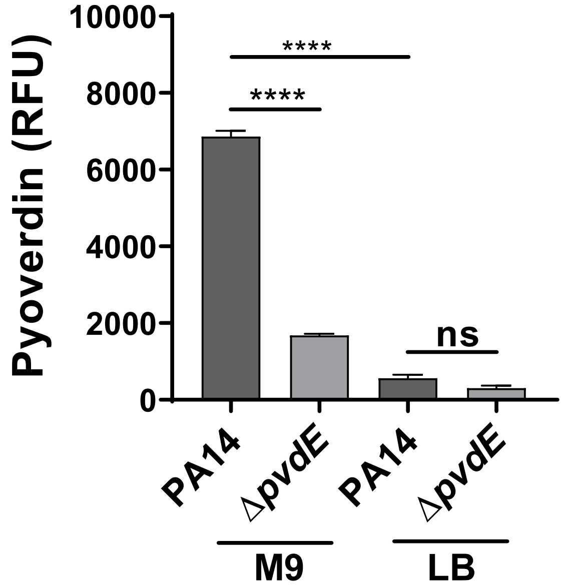


**Figure S5: Pyoverdine production is higher in M9 medium.** Pyoverdine was estimated by measuring fluorescence of cell-free supernatant after the growth in broth at 37°C for 24 hours. Fluorescence of blank medium is subtracted from test values for normalization. Δ*pvdE*, a pyoverdine production deficient mutant is used as control. An unpaired t test was used for analysis of significance (*, $P \leq 0.05$; **, $P \leq 0.01$; ***, $P \leq 0.001$).

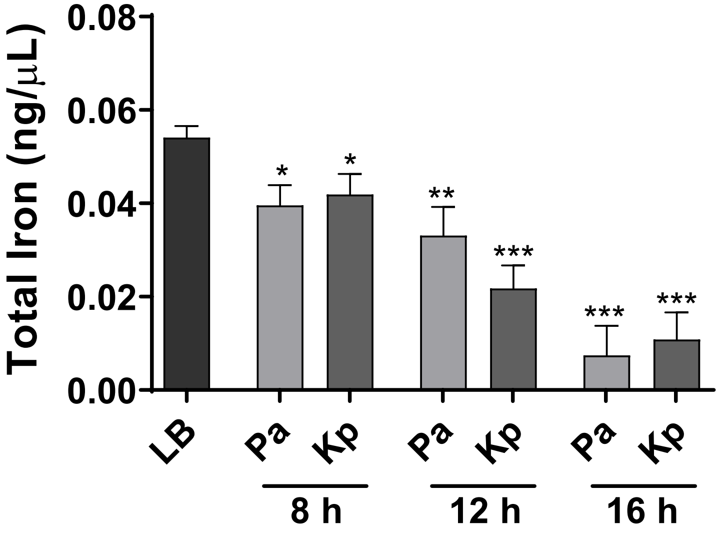


**Figure S6: Pa and Kp rapidly deplete iron from medium.** Total iron was estimated in the supernatant of Pa and Kp grown in LB at 8 h, 12 h and 16 h post inoculation. LB media was taken as control. An unpaired t test was used for analysis of significance (*, $P \leq 0.05$; **, $P \leq 0.01$; ***, $P \leq 0.001$).

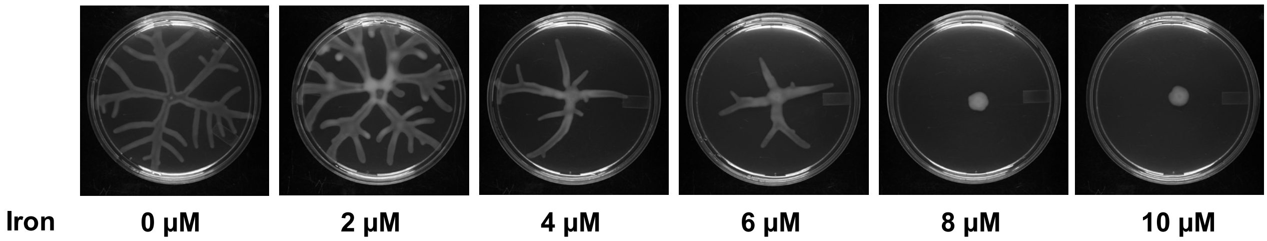


**Figure S7: Iron supplementation inhibits swarming in *P. aeruginosa*.** $FeSO_4.7H_2O$ is used for iron supplementation in M9 swarm agar in concentration from 0 to 10 µM. Swarm was allowed to develop for 24 hours at 37°C.
